# Implicit but not explicit exposure to threat conditioned stimulus prevents spontaneous recovery of threat potentiated startle responses in humans

**DOI:** 10.1101/304592

**Authors:** JP Oyarzún, E Càmara, S Kouider, L Fuentemilla, R de Diego-Balaguer

## Abstract

It has long been posited that threat learning operates and forms under an affective and a cognitive learning system that are supported by different brain circuits. A primary drawback in exposure-based therapies is the high rate of relapse when higher order inhibitory structures failed to inhibit the emotional responses driven by the defensive circuit. It has been shown that implicit exposure of fearful stimuli leads to a long-lasting reduction of avoidance behavior in patients with phobia through the facilitation of fear processing areas in the absence of subjective fear. Despite the potential benefits of this approach in the treatment of phobias and PTSD, implicit exposure to fearful stimuli is still under-investigated. Here, we used unconscious presentation of threat-conditioned stimuli in healthy humans, using a continuous flash suppression technique. We found that implicit exposure of a conditioned stimulus reduced, on the following day, defensive responses to the conditioned stimulus measured by threat-potentiated startle responses but not by the electrodermal activity. Our results suggest that implicit exposure using CFS might facilitate the modulation of the affective component of fearful memories, representing an important therapeutic target to further advance exposure-based psychotherapies.

## INTRODUCTION

The ability to learn that previously threatening stimuli are no longer a threat is critical for mental health since the disruption of this process can lead to anxiety disorders like phobias and post-traumatic stress disorder, PTSD. A long-standing critical issue in the treatment of threat-related memories is the high rate of relapse after initially successful therapy (Craske and Mystkowski, 2006).

It has been established that threat learning operates and forms supported by two distinct brain circuits (Hamm and Vaitl, 1996; LeDoux, 1993). The first is an affective learning system grounded in the defensive circuit based in the amygdala and operating implicitly (LeDoux, 1993). The second is a cognitive learning system, associated with the acquisition of the declarative knowledge of stimuli contingencies, expectancy of threat and conscious experience of fear that is sustained by hippocampal and prefrontal brain areas (Baeyens et al., 1995; Lang et al., 2000; LeDoux and Brown, 2017; Purkis and Lipp, 2001).

Exposure-based therapy is the most used procedure to treat threat-related memories (Rothbaum and Davis, 2003) and is founded on the principles of extinction learning (Craske, 1999; Milad and Quirk, 2012) where the threat-predicting stimulus (i.e., conditioned stimulus, CS) is repeatedly presented in the absence of the negative outcome (e.g., unconditioned stimulus, US). Through this procedure, subjects learn an inhibitory memory that, relying on prefrontal structures (e.g., dorsolateral and ventromedial prefrontal cortex) (Phelps et al., 2004; Schiller et al., 2013), suppresses the expression of the defensive responses initiated by amygdala-subcortical structures (Pare and Duvarci, 2012; Sotres-Bayon et al., 2006). However, this inhibitory function often fails, and defensive responses are spontaneously recovered with the passage of time (Rescorla, 2004).

It has been suggested that since extinction learning leaves the affective memory fairly intact (Baeyens et al., 1995; Myers and Davis, 2002), such implicit trace could later motivate fear recovery, especially when the inhibitory structures (i.e., the prefrontal cortex) are impaired, as is the case with anxiety-related patients (Konarski et al., 2007; Sotres-Bayon et al., 2006), or under stressful situations (J and Nadel, 1985). Some studies have shown that procedures that avoid prefrontal cortex (PFC) engagement to inhibit threat-related memories are highly effective in preventing the recovery of defensive responses to threat conditioned or phobic stimuli (Koizumi et al., 2016; Schiller et al., 2013; Siegel and Weinberger, 2009). Of particular interest are the works of Siegel and Weinberg (Siegel and Warren, 2013a, 2013b) showing that very brief repeated masked exposure to phobic stimuli led to long-lasting reduction in avoidance behaviour in spider-phobics. In a recent fMRI study, the authors (Siegel et al., 2017) suggested that the beneficial effects of masked exposure might have been mediated through a facilitation of threat memory processing and the activation of regulation areas as participants do not experience subjective distress during exposure. To date, however, implicit exposure is still under-investigated, harboring important theoretical as well as clinical implications.

Here, we investigated the effects of implicit exposure on a fearful memory after 24h, using a continuous flash suppression technique (CFS). To model fear acquisition and exposure-based therapy, healthy participants were threat conditioned on day 1 to fearful faces. On day 2 stimuli were presented through a stereoscope, either invisibly (through CFS) for the implicit group or explicitly for the explicit group. On day 3 participants were normally presented to threat conditioned stimuli and recovery of defensive responses were tested by analyzing threat-potentiated startle responses, electrodermal activity, and online expectancy reports.

As it has been suggested by other authors (“Anxious: The Modern Mind in the Age of Anxiety by Joseph E LeDoux, book review,” n.d.; Brewin, 2001; Siegel et al., 2017), we predict that by restraining cognitive-mediated fear processing, implicit exposure would promote threat memory processing at the implicit level and hinder the recovery of defensive responses.

## MATERIALS AND METHODS

### Participants

#### Implicit group

*Fifty-nine* (46 female, *M* = 22.95 years, *SD* = 3.78) healthy students with normal or corrected-to-normal vision were recruited for this group. On the first day, we excluded 16 participants that did not meet the threat acquisition criteria (see *exclusion criteria for acquisition*). From these, 23 more participants were excluded on the second day because they broke the suppression effect during implicit exposure (see *exclusion criteria for image suppression*). A final sample of 20 participants fulfilled the criteria for inclusion and followed the three consecutive-days experimental protocol of the implicit group.

#### Explicit group

Thirty-two healthy students with normal or corrected-to-normal vision were recruited for this group (25 female, *M* = 20.5 years, *SD* = 2.39). On day 1, we excluded 13 participants that were not threat conditioned and three that were non-responders (see *exclusion criteria for acquisition*). One participant did not return for day 3. A final sample of 15 participants fulfilled the criteria for inclusion and followed the three consecutive-days experimental protocol of the explicit group.

The study was approved by the Institute of Biomedical Research of Bellvitge ethics committee and all subjects from both groups signed an informed consent before their participation.

### Psychological Inventories

In order to control for psychological individual differences that could influence threat learning, all participants completed the Spanish version of the Spielberger State-Trait (STAI-T), the State-State (STAI-S) Anxiety Inventory (Spielberger, 1983) and the Spanish version of the 25-item English Resilience Scale (Wagnild and Young, 1993) containing the ‘Acceptance of Self and Life’ (ASL) and “Personal Competence” (PC) subscales.

### Stimuli

#### Visual Stimuli

We employed Ekman’s fearful faces (Ekman, 1976) as the conditioning stimuli (CS) as they can be processed in the absence of awareness through a rapid subcortical amygdala route (McFadyen et al., 2017). Faces were presented for 5 seconds with inter-trial intervals (ITI) of 10-12 seconds (after electrodermal activity was stabilized). Stimuli order presentation was randomized with the constraint that no more than 3 repetitions of the same stimuli occurred. Stimuli were displayed on a 22-inch computer monitor (resolution = 1,024 × 768 pixels; refresh rate = 60 Hz) and were controlled using Psychophysics Toolbox software (Brainard, 1997; Pelli, 1997). Stimulus contrast was equally set for all participants, at a level that was clearly visible when viewed on its own but was also easily suppressed with continuous flash suppression (CFS) (see CFS in experimental task below).

#### Electrical Stimulation

We used a mild electric shock to the wrist as the unconditioned stimulus (US) during threat conditioning on day 1. Shocks were delivered through an electrode attached with a Velcro strap to participants’ dominant inner wrist, with a maximum intensity of 15mA and 50 ms duration and co-terminated with faces presentation (Oyarzún et al., 2012). A Grass Medical Instruments stimulator (Grass S48 Square Pulse Stimulator) charged by a stabilized current was used with a Photoelectric Stimulus Isolation Unit (Model PSIU6). At the beginning of the session, participants regulated shock intensity to a level which they described as very uncomfortable yet not painful.

#### Air-puffs

In order to measure threat-potentiated startle responses (see below), we mechanically provoked blink responses by delivering 40 ms air-puffs, through a hosepipe directed to the anterior part of the temporal region between the outer canthus of the eye and the anterior margin of the auditory meatus (Haerich, 1994; Hawk and Cook, 1997) of the dominant-hand side. Airpuffs were delivered 4.5 s after every face presentation onset (did not overlap with electrical stimulations) and during every other intertrial interval (ITI). In order to habituate subjects to airpuff stimulation, each day started with 10 startle probes (Sevenster et al., 2012a).

### Experimental Task

#### Day 1. Fear acquisition

On day 1, participants were randomly presented with 3 fearful faces, 8 times each. Two of them (CS1 and CS2) co-terminated with a mild electric shock to the wrist on 75% of the trials (reinforcement was omitted in the 1st and 5th trial) and a third one was never followed by the aversive stimulus (neutral stimulus, NS). Face gender was counterbalanced and randomized across participants. To acquire asymptotic levels of learning, participants were instructed that two faces were going to be followed, most of the time, by an electric shock, while the third one was safe (Figure 1).

**Figure 1.**
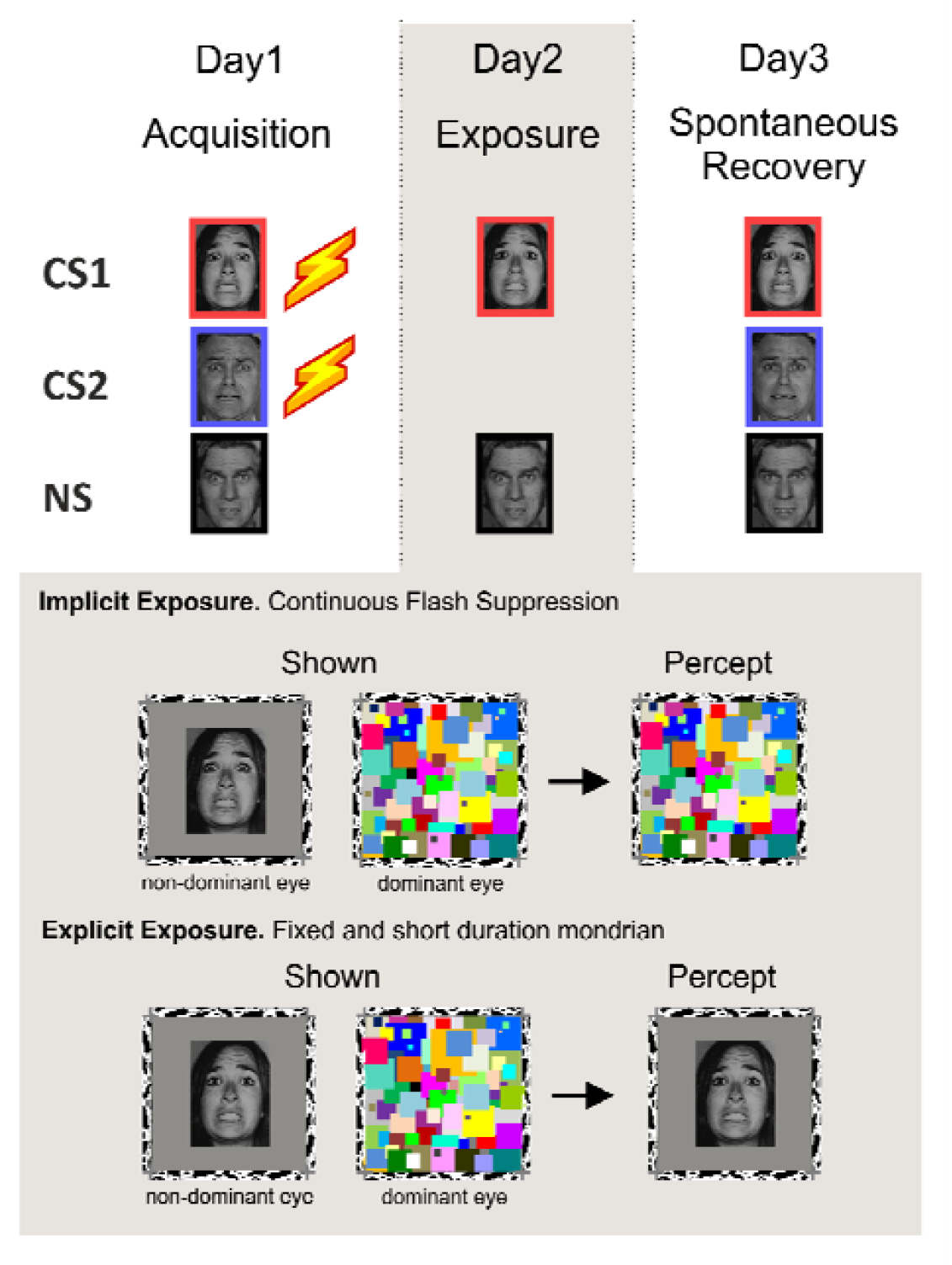
Three-day experimental design: acquisition exposure and spontaneous recovery test. Two faces were fear-conditioned on day 1 (CS1 and CS2) whereas a third face served as the neutral stimulus NS. CS1 and the NS were presented with no reinforcement on the second day using a continuous flash suppression (CFS) setting (with a stereoscope and colorful patches). These Mondrians were continuously flashing during picture presentation in the implicit group but were fixed and briefly presented in the explicit group. Acquisition on day 1 and Recovery Test on day 3 were conducted explicitly with faces at the center of the screen and without CFS setting.

##### Exclusion criteria for acquisition

The first exclusion criterion aimed to ensure that participants were fear-conditioned. We selected participants that showed differential electrodermal activity and startle potentiation to both threat-conditioned stimuli compared to neutral, this is; the average of the final 4 trials, in the acquisition session, for both CSs was greater than for the NS stimulus in the EDA or SR index. In addition, we excluded non-responder participants who showed below 0.02 μS peak to peak amplitude in the EDA index in more than 75% of unreinforced trials during acquisition (Raio et al., 2012).

#### Day 2. Exposure session

##### Implicit exposure group

Twenty-four hours after threat conditioning, using a stereoscope and the CFS technique (see CFS below) participants were unconsciously exposed with only two of the images presented on day 1: CS1 and NS, 16 times each in the absence of electric shocks. In order to control for participants’ awareness of the face presentation, we asked for a subjective report using the keyboard arrows. After each trial, they were asked: ‘Do you think you might have seen a face?’ ‘Yes’ or ‘No’, and then ‘Was it a male or a female?’ Subjects then indicated ‘male’ or ‘female’ and how sure they were of their answer with ‘sure’ or ‘not sure’.

###### Detection task

In order to dissuade participants to voluntary explore the non-dominant eye (by closing one eye) and thus break the CFS effect, we included a simple detection task on the dominant eye during Mondrian display (see CFS below). Three seconds after Mondrian onset, a central grey dot would randomly change to a different color for 1 second. At the end of the three awareness questions participants had to answer whether the dot had turned to green or not; although no feedback was received after each response participants were encouraged to be accurate in this task. Participants were pre-trained for this task in the training session (see training session below).

##### Exclusion criterion for image suppression

To ensure full image suppression we excluded participants that answered, in at least one trial: ‘yes’ to the first question (‘*Do you think you might have seen a face?*’) and were correct and confident (answered ‘sure’) in indicating the gender of the perceived face. Following this selection criteria, all the participants included in the final sample reported not seeing anything besides the Mondrian at all trials; this is, for every trial the participants included in the sample answered ‘*No*’ to the first question (except for one subject that answered seeing something on one trial), they all guessed faces at chance in the second question (main percentage of hits 46.71%; SD=7.5; [34.38% – 58.06%]) and responded to be ‘*not sure*’ about the guess in the third question. Participants learned to answer these 3 awareness questions during the training session on day 2.

##### Continuous Flash Suppression (CFS)

We employed the continuous flash suppression (CFS) technique, a binocular rivalry-based method capable of reliably suppressing visual awareness despite stimulus presentation for long periods of time (Fang and He, 2005; Lin and He, 2009; Tsuchiya and Koch, 2005). Using a mirror stereoscope (Stereoaids, Australia) placed 45 cm from the screen, we presented a continuously flashing colorful pattern (Mondrian) (at 10Hz) to the dominant eye and low-contrast (albeit visible) faces to the other, non-dominant eye. Mondrians were created with Matlab (MathWorks, Natick, MA) and the Psychtoolbox (Brainard, 1997; Pelli, 1997) and were presented for 5.5 seconds, starting 500 ms before face onset. In this manner, target faces were rendered invisible to the participants and thus processed without awareness. To determine eye dominance we used a sighting dominance test (Porac and Coren, 1976) where we asked participants to hold, with extended arms, a plastic board and look through a central small aperture to a picture placed on the wall at a 2-meter distance. The investigator would then cover one participants’ eye at a time and ask for a subjective report of the image. If the image was no longer seen when covering a certain eye then that eye was considered dominant.

###### Training session

After the eye dominance test and before starting the experiment on day 2, participants had a training session for 5 minutes to calibrate the stereoscope, ensure image suppression and familiarize participants with the task and questions. First, a black and white image of a zebra was presented to one eye and the zebra outline was presented to the other. The subjects adjusted the mirrors of the stereoscope using two knobs so that each eye in isolation saw either the full zebra or the full zebra outline, and with two eyes the zebra was aligned within the zebra outline. Then, subjects in both groups initiated a training sequence using 6 presentations of random objects (instead of the faces) where they were familiarized with the 3 awareness questions and, in the case of the implicit group, with the detection task.

##### Explicit exposure group

Participants followed the same procedure as the implicit group (same number, ITI, and length of stimuli presented through the stereoscope). However, for this group, face-pictures were explicitly presented. Mondrians were presented (in the dominant eye) for only 500 ms before face-picture presentation, so faces were fully visible to the participants, for the following 5 seconds (the same duration as in the implicit group), in the non-dominant eye. In the same way, as with the implicit group, the same three questions regarding picture awareness followed each image presentation. In order to encourage participants to pay attention to faces presentation, this group did not perform the color detection task. All participants reported seeing the faces at all trials; this is, they answered ‘*Yes*’ to the first question (except for one subject that reported not seeing a face on one trial), presented 100% accuracy in gender detection and were always sure about their response.

### Day 3

#### Spontaneous recovery test

After 24 hours we tested for recovery of defensive responses to all stimuli. Participants were presented with the three faces they saw on the first day, 6 times each in the absence of the shock. To remove attentional orienting effects on the first trials, an extra presentation of the neutral stimulus, which was not included in the analysis, was presented at the beginning of this session.

##### Online threat expectancy ratings

During the spontaneous recovery test participants had to indicate whether they expected to receive, or not, an electric shock after seeing each face on the screen. One second after face presentation the question ‘Are you expecting to receive a shock?’ appeared on the screen for 3 seconds. Participants answered, using the arrows of the keyboard, ‘Yes’, ‘No’ or ‘I don’t know’.

### Measures

#### Threat-potentiated startle responses (SR)

Startle responses were analyzed after delivery of air-puffs. We performed a monocular electromyography (EMG) on the orbicularis ocular muscle of the dominant eye. A 6 mm Ag/AgCl electrode filled with a conductive gel was placed 1.5 cm below the lower eyelid in line with the pupil at forward gaze, a second electrode was placed 2 cm lateral to the first one (center-to-center), and a signal ground electrode was placed on the forehead 2 cm below the hairline (Blumenthal et al., 2005).

##### EMG data analysis for SR

Raw EMG data were notched and band-pass filtered (28-500 Hz, Butterworth, 4^th^ order), and afterward rectified (converting data points into absolute values) and smoothed (low-pass filter 40 Hz) (Blumenthal et al., 2005). Peak blink amplitude was determined in a 30-150 ms interval following air-puff delivery. EMG values were standardized using within-participant Z scores for each day, and outliers (Z > 3) were replaced by a linear trend at point (Sevenster et al., 2012a). For comparisons between exposure on day 2 and spontaneous recovery test on day 3, Z scores were calculated using both exposure and recovery test data. For comparisons within stimuli (CS1, CS2 and NS) on day 3, Z scores were calculated using only recovery test data.

#### Electrodermal activity (EDA)

Electrodermal activity and EMG was sampled at 1000Hz and was recorded during the whole session using Brain Amps amplifiers. EDA was assessed using two Ag-AgCl electrodes connected to a BrainVision amplifier. The electrodes were attached to the middle and index fingers of the non-dominant hand.

##### EDA data analysis

EDA waveforms were low-pass filtered (1Hz) and analyzed offline with Matlab 7.7. F. Single-trial changes in EDA were determined by taking the base-to-peak difference for a 4.5 s window after stimulus onset and before air-puff (or electric shock) delivery. The resulting amplitude of the skin conductance response (SCR) value was standardized using within-participant Z scores for each day, and outliers (Z > 3) were replaced by a linear trend at point (Sevenster et al., 2012a). As for EMG analyses, comparisons between exposure on day 2 and spontaneous recovery test on day 3 used Z scores calculated using both exposure and recovery test data. For comparisons within stimuli (CS1, CS2 and NS) on day 3, Z scores were calculated using only recovery test data.

### Online US-expectancy ratings (OER)

Since explicit evaluation of contingencies could affect learning during fear acquisition and extinction learning during exposure, expectancy ratings were made only during day 3. After each image presentation, the question “Are you expecting to receive an electrical shock?” appeared on the top of the screen for 3.5 seconds to which participants answered “*Yes*” (scored 3), “*No*” (scored 1) or “*I don’t know*” (scored 2) using the keyboard. Participants were encouraged to maintain their hands over the keyboard at all times and to restrict hand and head movement as much as possible.

## RESULTS

### Acquisition

#### Equivalent levels of threat acquisition for conditioned stimuli in both groups and in both measures

##### Threat Potentiated Startle responses (SR)

A two-way mixed analyses of variance (ANOVA) with group (implicit versus explicit) as a between-subject factor and stimuli (CS1 CS2 and NS) as a within-subject factor showed equivalent levels of SR for both groups in the last 4 trials (all p values > .1 for group and group x stimuli interaction) but a main effect of stimuli (F_(2,66)_ = 12.23; *p* < .001; 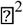 = .27) (Figure 2A). A Repeated Measures ANOVA (RM-ANOVA) combining both groups showed successful threat conditioning results: a main effect of stimuli (F_(2,68)_ = 12.12; *p* < .001; 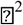 = .26) with equal responses for CS1 and CS2 (Paired t-test, *t*_34_ = -.59; p = .55; *d* = .10) that were greater in comparison with NS (Paired t-testCS1-NS, *t*_34_ = 4.51; *p* < .001; *d* = .76, CS2-NS *t*_34_ = 3.75; *p* = .001; *d* = .63).

**Figure 2.**
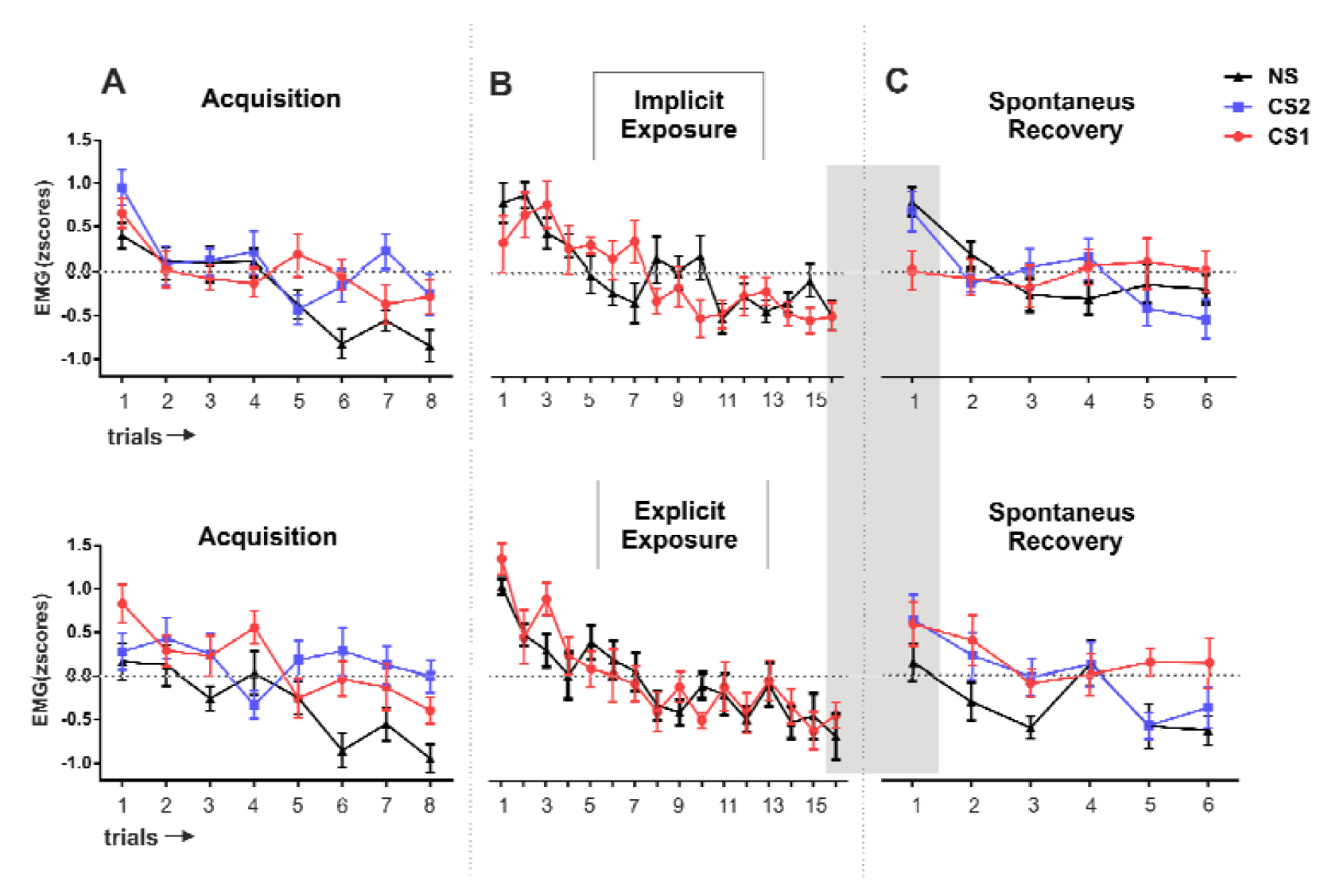
Threat potentiated startle responses trial by trial throughout the 3-day experiment for both experimental groups. The top panel depicts implicit exposure group. The lower panel depicts explicit exposure group. Mean standardized startle responses were calculated using all trials within each session for each group. Plots represent the mean response of (A) CS1 CS2 and NS during acquisition on day 1 (B) CS1 and NS during exposure on day 2 (C) CS1 CS2 and NS during spontaneous recovery on day 3. EMG: electromyography. Error bars represent standard error of the mean (SEM).

##### Electrodermal activity (EDA)

EDA analyses showed similar results. Responses were equivalent between groups (all p values > .1 for group and group x stimulus interaction) but a main effect of stimuli was observed (F_(2,66)_ = 26.61; p < .001; 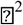 = .44) (Figure 3A). A RM-ANOVA combining both groups showed successful threat conditioning results with a main effect of stimulus (F_(2,68)_ = 28.48; *p* < .001; 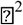 =.45) where CS1 and CS2 showed equivalent responses (paired t-test *t*_34_ =.29; *p* = .76; *d* = .05) but greater than the NS (CS1-NS *t*_34_ = 6,04; *p* < .001; *d* = 1.02, CS2-NS *t*_34_ = 5.88; *p*< .001; *d* = .99).

**Figure 3.**
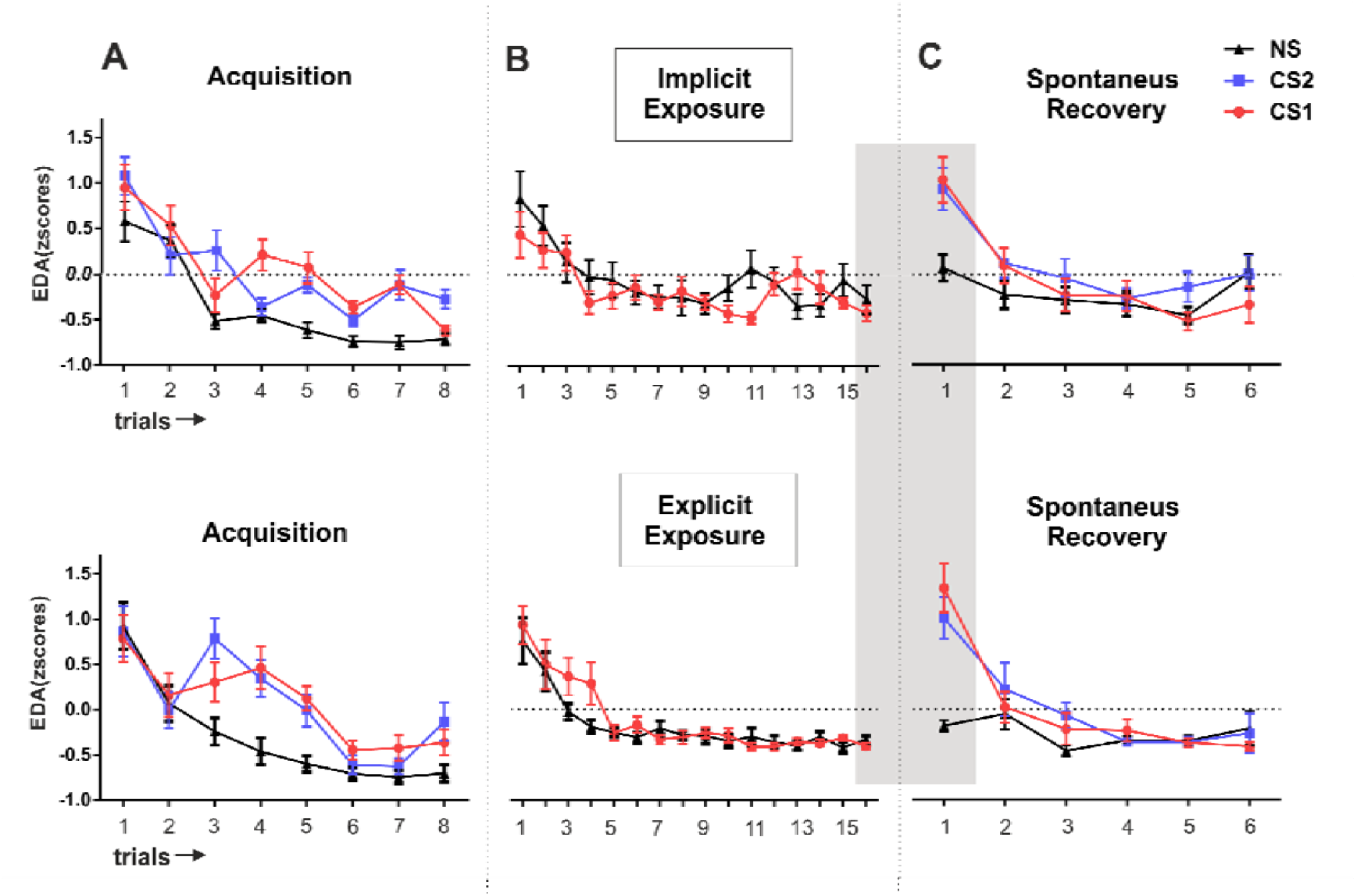
Trial-by-trial electrodermal activity throughout the 3-day experiment for both experimental groups. The top panel depicts implicit exposure group. The lower panel depicts explicit exposure group. Mean standardized electrodermal activity was quantified using all trials within each session for each group. Plots represent the mean electrodermal activity of (A) CS1 CS2 and NS during acquisition on day 1 (B) CS1 and NS during exposure on day 2 (C) CS1 CS2 and NS during spontaneous recovery on day 3. EDA: electrodermal activity. Error bars represent standard error of the mean (SEM).

### Exposure session

#### Gradual overall decrease of responses during exposure session with no differences between groups nor between stimuli, in both measures

We then analyzed the course of extinction learning during exposure using a two-way mixed ANOVA with group (implicit versus explicit) as an inter-subject factor and stimulus (CS1 and NS) and time (first trials 1-2 and last trials 15-16) as intra-subject factors.

##### Threat Potentiated Startle reflex (SR)

We found no differences in responses between groups nor differential responding between stimuli (all *p* values > .5 for group, stimulus, and group x stimulus interaction). When looking at differences across time we found a decrease in responses from beginning to end of the session (main effect of time; F_(1,33)_ = 55.57; *p* < .001; 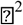 = .62) that was equivalent between groups and stimuli (all *p* values > .1) (Figure 2B).

##### Electrodermal activity (EDA)

EDA analyses showed similar results; no differences between groups nor between stimuli (all *p* values > .1 for group, stimulus, and group x stimulus interaction) (Figure 3B). Again, we found a decrease in responses from beginning to end of the session (main effect of time; F_(1,33)_ = 57.50; *p* < .001; 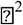 = .63) that was equivalent between groups and stimuli (all *p* values > .1).

### Spontaneous Recovery Test

To test the recovery of defensive responses on day 3, we compared the last trial of the exposure session with the first trial of the spontaneous recovery test for CS1 and NS (Oyarzún et al., 2012; Schiller et al., 2010, 2013; Soeter and Kindt, 2011; Warren et al., 2014) (Figure 4).

**Figure 4.**
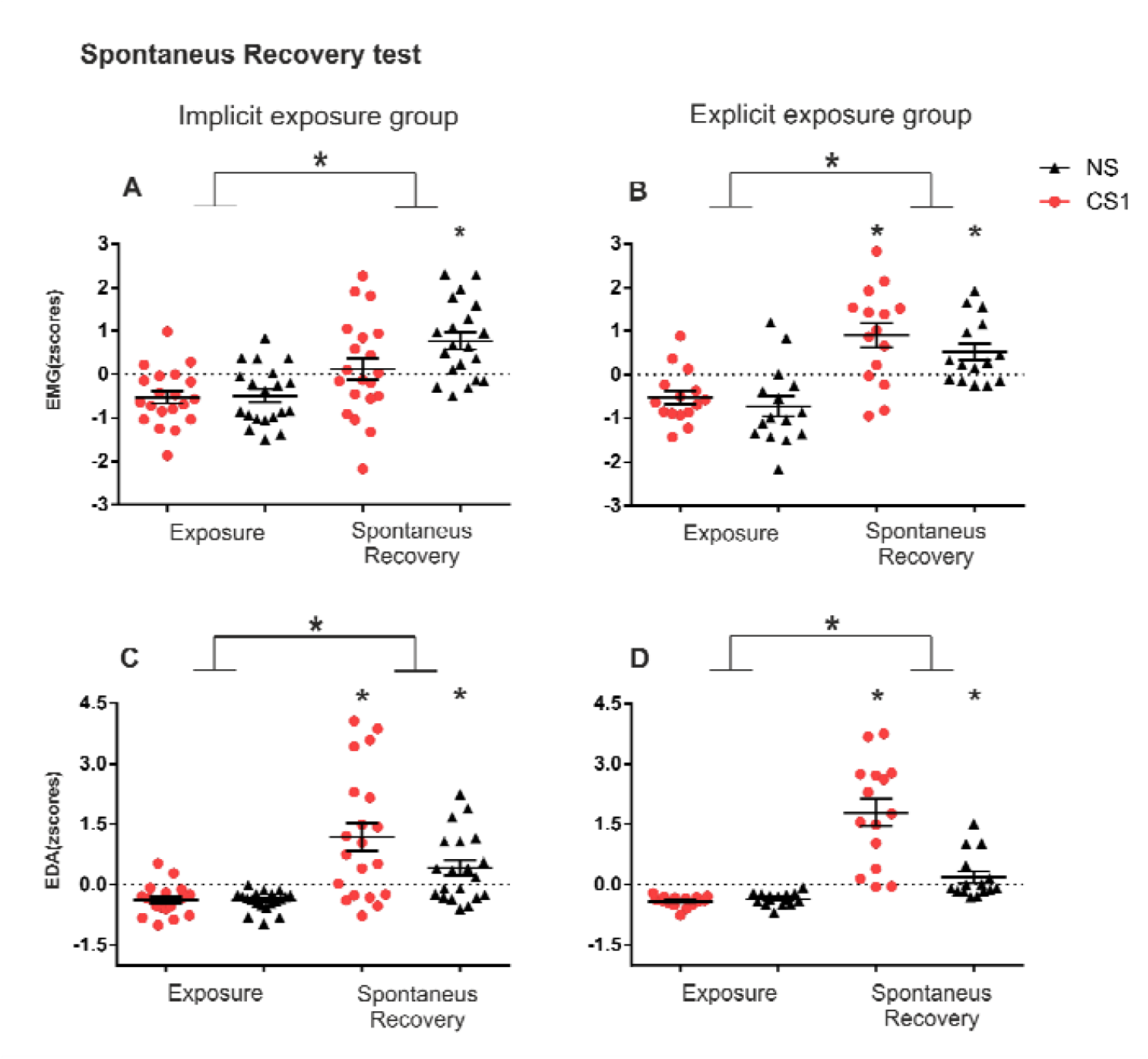
Recovery of defensive responses for both experimental groups. Mean standardized startle responses were calculated using all trials for CS1 and NS during exposure and spontaneous recovery sessions for each group in each measure. Plots represent the last trial of exposure on day 2 and the first trial of spontaneous recovery on day 3. (A) Mean of startle response for the implicit group. (B) Mean of startle response for the explicit group (C) Mean of electrodermal activity for the implicit group. (D) Mean of electrodermal activity for the explicit group. EMG: electromyography; EDA: electrodermal activity, small *: p < .05 comparison for each stimuli between phases, big*: main effect of phase. Error bars represent standard error of the mean (SEM).

We first compared recovery of defensive responses between groups, for the startle responses and for the electrodermal activity. And secondly, we compared the differential responses between the EDA and the SR measures within each group.

#### CS1, in the implicit group, showed no recovery of responses from the end of exposure session to the beginning of the recovery test. Only CS1 in the implicit group showed lower responses in comparison with CS2

##### Threat potentiated Startle Reflex (SR)

A two-way mixed ANOVA with group (implicit versus explicit) as a between-subjects factor, and phase (exposure and recovery test) and stimulus (CS1 and NS) as within-subject factors, revealed no main effect of group (F_(1,33)_ = .30, *p* = .58; 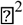 = .00). A main effect of phase (F_(1,33)_ = 38.92; *p* < .001; 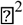 = .54) that was equivalent between groups (phase x group _(1,33)_ = 1.06; *p* = .31; 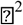 = .03) indicated that SR responses increased at recovery in both groups. However, we found a significant stimuli x group interaction (F_(1,33)_ = 10.0098, *p* < .005; 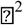 = .23) (Figure 4A-B). We thus compared stimuli responses between groups. Unpaired t-test showed similar responses for the NS in both groups (*t*_(33)_ = 1.52; *p* = 1.13; *d* = .49) but lower responses for the CS1 in the implicit than the explicit group (t_(33)_ = −2.19; *p* = .03; *d* = .74). Intra-group comparison of stimuli showed, in the implicit group, lower responses for the CS1 in comparison with the NS (*t*_(19)_ = −2.97; *p* = .008; *d* = .66). In contrast, similar responses for NS and CS1 were found in the explicit group (*t*_(14)_ = 1.68; *p* = .11; *d* = .43), indicating that implicit but not explicit exposure reduced SR responses to CS1.

We then compared CS1 responses with CS2 on day 3; another homologous stimulus that was equally threat conditioned in the first session, but that was not exposed to participants on day 2 (Figure 2C). A two-way mixed ANOVA with group (implicit versus explicit) and stimulus (CS1, CS2 and NS, standardized within day 3) as a between and within-subject factors respectively, revealed a significant group x stimulus interaction (F_(2,66)_ = 3.93; p = .02; 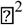 =.11). Whereas in the explicit group all stimuli (i.e. CS1 NS CS2) showed comparable high responses (all p values > .1), differences across stimuli were found in the implicit group (implicit F_(2,38)_ = 3.44; p = .04; 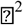 = .15, explicit F_(2,28)_ = 1.37; p = .26; 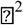 = .09), where only CS1 showed reduced response compared to the CS2 (*t*_(19)_ = −2.09; *p* = .04, *d* = .46) and NS (*t*_(19)_ = −2.77; *p* = .01, *d* = .62).

#### EDA remained equivalent in both groups, with an overall increased activity from exposure session to recovery test but greater recovery for CS1s that was comparable to CS2s responses

##### Electrodermal activity (EDA)

A two-way mixed ANOVA with group (explicit versus implicit) as an inter-subject factor, and phase (exposure and test) and stimuli (CS1 and NS) as within factors revealed a main effect of phase (F_(1,33)_ = 70.04; p < .001; 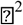 = .69), stimuli (F_(1,33)_ = 15.98; p < .001; 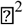 = .32) and phase x stimuli interaction (F_(1,33)_ = 18.02; p < .001; 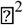 = .35), but no differences were found between groups (all *p* values > .1 for group, group x stimulus and, group x stimulus x phase interaction) (Figure 4 C-D). We thus combined groups and compared stimuli responses between phases. As expected, responses significantly increased from the end of the exposure session to the recovery test in both stimuli (paired t-test NS *t*_(34)_ = −5.37; *p* < .001; *d* = −0.90, CS1 *t*_(34)_ = −7.10; *p* < .001, *d* = −1.2). And, although responses between stimuli were comparable at the end of the exposure session (*t*_(34)_ = -.40; *p* = .68; *d* = -.06), responses in the recovery test were greater for CS1 than for NS (*t*_(34)_ = 3.93; *p* < .001; *d* = 0.66). Thus, showing that in both groups, CS1 and NS, incremented EDA responses from the end of day 2 to test, but with greater recovery for CS1.

We then explored whether such recovery in the CS1 was similar to the response of its conditioned homologous CS2 on day 3 (Figure 3C). A mixed ANOVA with group and stimuli (CS1, CS2 and NS) showed no differences across groups (all *p* values > .5 for group and group x stimulus interaction) but a main effect of stimulus (F_(2,66)_ = 15.21; *p* < .001; 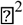 = .32) that was driven by equal responses for CS1 and CS2 on day 3 (paired t-test *t*_(34)_ = .70, *p* = .48, *d* = .11) but greater than NS (CS1-NS *t*_(34)_ = 5.48, *p* < .001, *d* = .92, CS2-NS *t*_(34)_ = 5.15, *p* < .001, *d* = .87). Thus, in the EDA measure, regardless of type of exposure, conditioned stimuli CS1 showed equivalent increased recovery than CS2 on day 3.

#### Comparisons between measures within groups. The implicit group showed a down-modulation of CS1 in the recovery test in the SR but not in the EDA. In contrast, the explicit group showed greater responses for CS1 and CS2 than NS in both the EDA and SR

In order to directly compare the differential responses in the EDA and the SR measures we tested whether the recovery of defensive responses was different between measures within each group. We performed RM-ANOVA with measure (SR and EDA), phase (exposure and test) and stimulus (CS1 NS) as within-subject factor, separately for each group. The implicit group showed a main effect of stimuli (F_(1,19)_ = 44.83, p < .001; 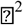 = .70), and interestingly a significant interaction of stimulus x measure (F_(1,19)_ = 8.81, p < .01; 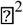 = .31) and stimulus x measure x phase (F_(1,19)_ = 6.32, p = .02; 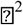 = .25) (Figure 4 A-C). Follow-up paired t-test showed greater CS1 responses from exposure to test in the EDA measure (t_(19)_ = −4.15, p < .005; d = -.92) but not in the SR measure (t_(19)_ = −1.87, p = .07; d = -.42). In contrast, the NS showed increment of responses in both measures (EDA t_(19)_ = −4.13, p = .001; d = -.92; SR t_(19)_ = −4.2, p < .001; d = -.95). Thus, indicating that CS1 responses were divergently down-modulated in the SR index but not in the EDA.

The explicit group showed a main effect of phase (F_(1,14)_ = 66.65, p < .001; 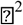 = .82), stimuli (F_(1,14)_ = 10.72, p = .006; 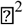 = .43) and again significant interactions of: stimulus x phase (F_(1,14)_ = 8.32, p = .01; 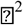 = .37), stimulus x measure (F_(1,14)_ = 7.66, p = .01; 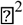 =.35) and stimulus x phase x measure (F_(1,14)_ = 5.25, p = .03; 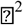 = .27) (Figure 4 B-D). First, we looked for stimuli responses increments from the exposure session to test. In this case, significant increments were found in both measures for both stimuli [CS1 (EDA *t*_(19)_ = −6.77, *p* < .001; *d* = −1.51; SR *t*_(19)_ = −4.01, *p* = .001; *d* = -.89), and NS (EDA *t*_(14)_ = −3.59, *p* = .003; *d* = -.92, SR *t*_(19)_ = −3.74, *p* = .002; *d* = -.96). We then looked for stimuli responses between phases. In the recovery phase, we found that whereas CS1 and NS responses were equivalent in SR (paired-t-test *t*_(14)_ = −1.34, *p* = .20, *d* = -.34) responses in the EDA were greater for the CS1 (paired-t-test *t*_(14)_ = 4.25, *p* = .001, *d* = 1.09). Thus showing that the explicit group increased responses for both stimuli in both measures but in the EDA the recovery was greater for the CS1.

### Online Threat Expectancy Ratings (OER) on day 3

#### Participants’ explicit contingency learning was not modulated by either implicit or explicit exposure

We then explored on day 3 whether participants expected to be shocked after the presentation of the faces (Figure 5). A two-way mixed ANOVA with group (implicit versus explicit) as between-subject factor and stimuli (CS1, CS2 and NS) and time (mean of the first 2 trials versus mean of the last 2 trials) as within-subject factor showed no differences between groups (all p values > .1 for group, group x stimuli and group x time interaction). Thus, these results indicated that our experimental manipulation did not affect OER. However we found a main effect of stimuli (F_(2,66)_= 50.34; *p* < .001, 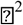 = .33), time (F_(2,66)_ = 16.25; *p* < .001, 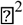 = .33) and stimuli x time interaction (F_(2,66)_ = 5.16; *p* < .005, 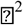 = .13). We thus explored stimuli responses across time. We found that participants expectancy scored for CS1 and CS2 stimuli decreased from beginning to the end during the recovery session (CS1 *t*_(34)_ = 3.39, *p* < .005; *d* = .57, CS2 *t*_(34)_ = 3.72, p < .005; d = .63). Interestingly and congruent with the threat generalization responses to the NS in the first trials of the recovery test, NS also showed a decrease of responses from beginning to the end of session (NS *t*_(34)_ = 2.71, *p* = 01; *d* = .45), as some subjects reported not to be sure of expecting to be shocked when presented with the NS (scored = 2) in the first trials. As expected, although shock expectancy was similar between CS1 and CS2 at both the beginning and end of the session (all p values > .5), NS scores were significantly lower at both the beginning (CS1-NS *t*_(34)_ = 8.90, *p* < .001; *d* = 1.52, CS2-NS *t*_(34)_ = 8.44, *p* < .001; *d* = 1.42) and end of the session (CS1-NS *t*_(34)_ = 5.60, *p* < .001; *d* = .94, CS2-NS *t*_(34)_ = 5.23, *p* < .001; *d* = .88). Thus, participants maintained the cognitive threatful representation for conditioned stimuli from the beginning to the end of the session.

**Figure 5.**
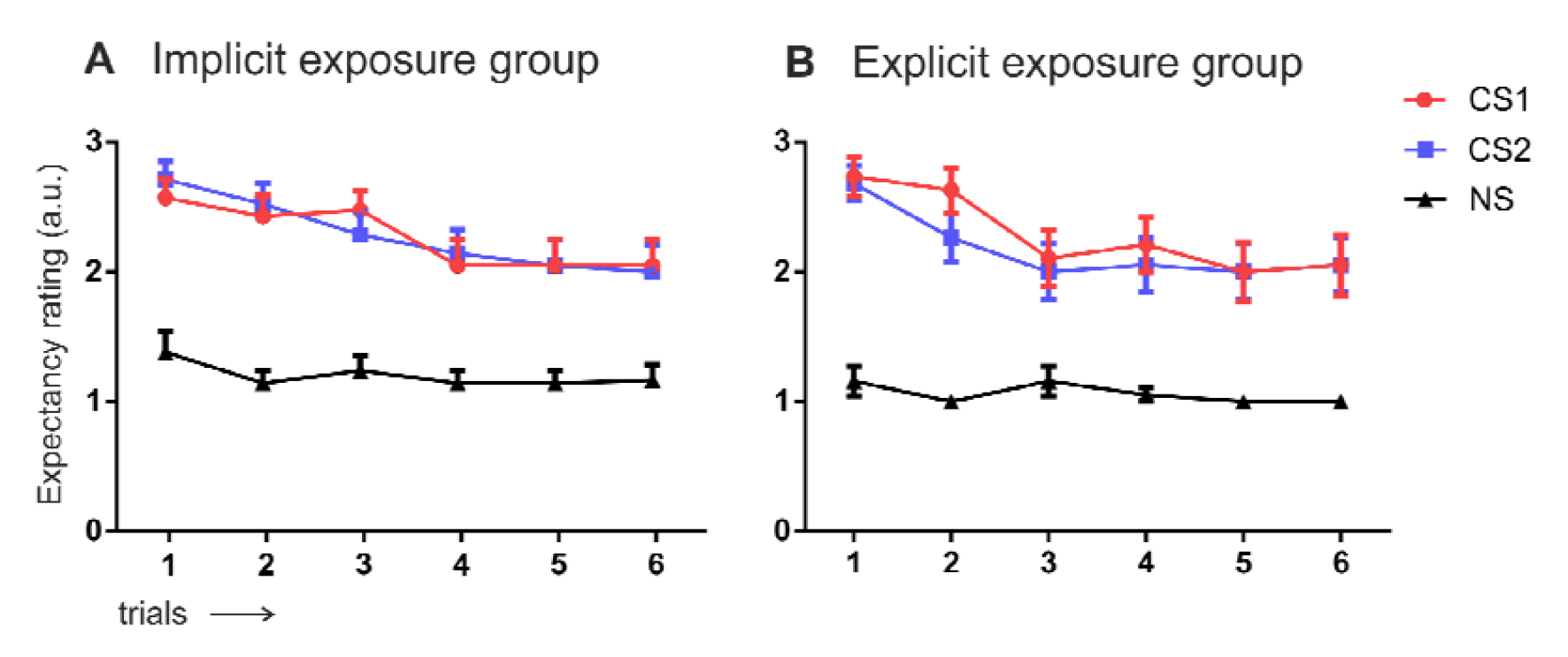
Online threat expectancy ratings during recovery test. During each picture presentation, subjects indicated whether they either expected (pressed 3), did not expect (pressed 1) or were not sure about (pressed 2) imminent shock occurrence. Error bars represent standard error of the mean (SEM); (a.u) arbitrary unit.

## PSYCHOLOGICAL INVENTORIES

### Equivalent scores between groups

Since anxiety traits have been previously related to aspects of implicit emotional learning (Raio et al. 2013) we checked whether our participants presented equivalent scores between groups in the psychological inventories. No significant differences were found between groups in any of the psychological inventories (see Table 1 for descriptive statistics); participants showed similar scores in the Spanish version of the STAI-state Inventory (unpaired t-test; *t*_(33)_ = -.55, *p* = .58, *d* = .19), the STAI-trait inventory (*t*_(33)_ = −1.55, *p* = .12, *d* = .52) and the Spanish version of the 25-item English Resilience with ASL and PC subscales (Group, F_(1,33)_ = .69; *p* = .41; 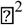 = .02, group x scale interaction, F_(1,33)_ = 1.00; *p* = .32; 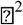 = .03).

**Table 1.** Descriptive statistics of inventory scores

| Inventory | Implicit |  | Explicit |  |
| --- | --- | --- | --- | --- |
|  | Mean | SD | Mean | SD |
| **STAI-state** | 10.15 | 5.33 | 11.06 | 4.13 |
| **STAI-trait** | 9.7 | 7.6 | 13.0 | 3.4 |
| **PC** | 91.22 | 12.56 | 91.46 | 9.04 |
| **ASL** | 37.11 | 5.77 | 40.26 | 4.11 |

*Note:* PC = Personal Competence; ASL = Acceptance of Self and Life

*Note:* PC = Personal Competence; ASL = Acceptance of Self and Life

These results indicate that the differences observed for the implicit and explicit groups are unlikely to be due to differences in anxiety and resilience traits between the groups.

## DISCUSSION

Two groups of participants underwent a partial reinforced threat-conditioning paradigm using three fearful faces. Two of the faces co-terminated with a mild electric shock to the wrist on 75% of trials (conditioned stimuli; CS1 and CS2) while a third face served as the neutral stimulus (NS).

On the second day, one group of participants underwent implicit while the other underwent explicit exposure to one of the threat conditioned stimulus. For the implicit condition, CS1 and NS were presented unconsciously using the continuous flash suppression (CFS) technique and no shocks were administered, while CS2 was not presented. The explicit group followed the same procedure except that pictures were explicitly presented (see Materials and Methods section). On the following day, we tested spontaneous recovery, by presenting all participants explicitly with the three faces in the absence of electric shocks (see design in Figure 1). We used a combination of measures to examine defensive responses: threat-potentiated startle reflex (SR), electrodermal activity (EDA) and online expectancy ratings (OER).

We found that exposing participants implicitly with previously threat conditioned stimulus reduced the recovery of defensive responses after 24h, measured by SR, but not by EDA or OER.

Our results highlight the divergent expression between two physiological measures (EDA and SR) where implicit exposure only modulated threat potentiated SR. Dissociation between both measures has long been recognized and although there is still much debate about the nature of each measure it has been suggested that they are differently modulated by different neural systems during threat memory encoding, extinction, and retrieval (Sevenster et al., 2014; Soeter and Kindt, 2010).

In our experiment on day 3, EDA followed a similar pattern of responses as those presented by the OER, but only at the beginning of the test session; higher responses for CS1 and CS2 than for NS that gradually decreased throughout the session. Such correspondence across both measures fits well with the idea that EDA is sensitive to modulations of threat explicit expectancies (Lovibond, 2003; Sevenster et al., 2014; Soeter and Kindt, 2010). However, the fact that OER and EDA, increasingly dissociated as the session progressed; with a stronger drop in EDA to all stimuli (Figure 3) but sustained high OER (Figure 5) suggest that EDA reflects subjective feelings of fear and it might behave independently from contingency knowledge, as reported in other studies (Raio et al., 2012). Indeed, previous studies suggest that EDA is driven by amygdala activity (Koizumi et al., 2016; Schiller et al., 2013).

Critically, the fact that implicit exposure only modulated SR in the first trial during the recovery test might suggest that SR is more sensible than EDA, to subtle modulations in the affective system, potentially induced during implicit exposure of CS1. In fact, SR as an automatic reflex, has been considered to be tightly regulated by the defensive circuit reflecting amygdala activity for negative affective valence (Hamm and Vaitl, 1996; Lang et al., 1990), whereas EDA appears be more sensible to cognitive modulations by the explicit expectations of upcoming relevant events (Sevenster et al., 2012b, 2014). Critically, if this is the case, our results would suggest that implicit exposure might separately modulate the implicit trace of fearful memories.

Of note, NS showed an increment of defensive responses in the recovery test in both groups and for both measures (when comparing the last trial of the exposure session with the first trial of the spontaneous recovery test session), suggesting a global threat generalization effect. Generalization in the physiological responses was further supported by the results in the OER where participants reported to be ‘not sure’ of being shocked with NS presentation in the first trials on day 3. Generalization of defensive responses in this type of paradigm has been reported previously by other studies (Kindt and Soeter, 2013; Oyarzún et al., 2012; Soeter and Kindt, 2011). In the context of our current design, it is possible that threat generalization was transferred via shared element among all stimuli (Dunsmoor and Murphy, 2015); this is, airpuffs which were always presented at the end of each picture (to induce the blinking response) (see Materials and Methods section), and were frequently followed by the electric shock (75% of times for the CSs).

An important point to consider is the fact that no differential responses between conditioned and neutral stimuli, nor between groups (implicit vs explicit) were observed throughout the course of the exposure session. One possible explanation is that the use of the stereoscope during exposure (and not during day 1 or 3) added new contextual cues that impaired the retrieval of threatful associations and precluded discrimination among stimuli. The use of the stereoscope only on day 2 was aimed to increase ecological validity of the exposure task, as the acquisition of fear associations and re-exposure to fearful stimuli would be unlikely to occur throughout a stereoscope in a real context.

Our results are consistent with and build on previous studies using a very brief exposure (VBE) approach, in which pictures of spiders were presented very rapidly (i.e., 25 ms) in phobic patients, leading to long-lasting reduction of avoidance behavior (Siegel and Warren, 2013a, 2013b). In an attempt to look for the mechanism underlying this effect, the authors (Siegel et al., 2017) scanned patients while exposed to either masked or clear visible phobic stimuli (in two separated groups). Counterintuitively, they showed that presentations of either masked or visible phobic stimuli activated or deactivated, respectively, brain regions that support emotional regulation like ventromedial PFC. They posited that limited awareness during exposure and lack of subjective fear as well as amygdala activity reduction might facilitate fear processing and emotional regulation. In addition, in other studies, it has been shown that when the prefrontal cortex is not engaged during extinction learning (due to a lesion or due to early development stage) subjects do not present recovery of defensive responses and amygdala is more involved during extinction, leading to a permanent extinction (Kim and Richardson, 2010; Koenigs et al., 2008). These results, point out the possibility that implicit exposure in our experiment might have engaged similar mechanism that leads to attenuation of defensives responses, albeit only detected by SR measure.

Although the neural mechanism underlying CFS suppression effects are still largely unknown, a functional neuroimaging study using CFS and invisible presentations of fearful faces (Lapate et al., 2016) showed that while awareness of cues promoted PFC-amygdala functional connectivity, invisible presentation of faces did not engage such regulatory circuit. In the case of our implicit exposure paradigm, it is possible that faces are repetitively processed by the amygdala, via a fast subcortical pathway (Méndez-Bértolo et al., 2016) and by sensory areas representing CS while unaware and thus in the absence of activation of the defensive circuit. This, in turn, would promote emotional memory processing perhaps by the desensitization of low-level threat related regions, as posited by Siegel et al. 2017 (Siegel et al., 2017). In fact, it has been reported that ex-spider phobic patients that showed permanent extinction after 6 months presented low activity in ventro-visual regions that were hyperresponsive to spiders before the therapy. Further, the authors revealed that reduced activity in a restricted portion of the same visual cortical region (right lateralized lingual gyrus) immediately after therapy predicted long-term permanence of extinction learning (Hauner et al., 2012). These results suggest that tapping into sensorial and low-level defensive networks might change the association between stimulus and defensive response, leading to permanent extinction without the need of prefrontal inhibitory control. As our experiment cannot account for any neural mechanism underlying CFS exposure further research is still imperative to examine such argument.

An important disadvantage and methodological limitation from CFS technique is that image suppression is often broken when presenting threatful images (Yang et al., 2007) which might limit its use as a sole tool in clinical settings. In our experiment, to reduce subject attrition by image suppression failure, we implemented a task where participants had to report the color of a central dot within the Mondrian. However, the fact that participants needed to hold their answer for a couple of seconds, might have comprised higher cognitive demand load during image presentation. Whether this could have affected our results is unknown and more research would be needed to clear out this possibility. Despite implementation of this task, around half of our participants needed to be ruled out in this study for having broken the suppression effect. The fact that the selection criteria eliminated so many participants might constitute a potential confound as selected participants might share psychological features that make them more likely to show reduced defensive responses in the SR during spontaneous recovery. Although our selected participants rated equivalent scores in all psychological inventories, these results urge the need for further investigation and replications that could circumvent bias selection of participants by improving suppression effect during CFS exposure.

Our results deviate from those of Golkar & Ohman (2012) (Golkar and Öhman, 2012). In their experiment, the authors extinguished two conditioned stimuli; one under masked and the other under visible conditions. In contrast to our results, in their study the stimulus that was unconsciously extinguished presented more fear recovery than the one explicitly extinguished. However, it might well be the case that the parallel engagement of explicit and implicit learning could jeopardize the latter, as both explicit and implicit systems share encoding resources (Turk-Browne et al., 2006, p.) and might interact in a competitive manner (Kim and Baxter, 2001). Indeed, it has been suggested that the strong cognitive component of exposure-based therapies may actually preclude extinction learning at the implicit level (“Anxious: The Modern Mind in the Age of Anxiety by Joseph E LeDoux, book review,” 2015).

We believe that implicit exposure using CFS might promote processing of fearful memories in the subcortical threat related networks and facilitate emotional regulatory areas. The fact that fearful stimuli are experienced in the absence emotional distress in patients might help to change the threatful trace and improve the course of the therapy. Although our results provide encouraging evidence supporting these ideas, our findings call out the need for further investigation to circumvent methodological limitations, precise the mechanism involved, and uncover the potential of CFS implicit exposure as a valuable complementary procedure to further advance exposure-based psychotherapies.

## Acknowledgements

The authors would like to thank Bert Molenkamp and David Cucurell for their technical assistance and Dieuwke Sevenster and Joaquin Moris for their advice on the experimental methodology. They are also thankful to Miguel Fullana and Rafael Torrubia for supplying critical equipment for this study and Ivonne Heideman for helping during data collection.

## Author Contributions

J.O. conducted the experiments and analyzed the data, J.O., R.D.B and L.F. designed the experiments and wrote the paper, and J.O., E.C., and S.K. programmed the task.

## REFERENCES

Baeyens, F., Eelen, P., Crombez, G., 1995. Pavlovian associations are forever: On classical conditioning and extinction. J. Psychophysiol. 9, 127–141.

Blumenthal, T.D., Cuthbert, B.N., Filion, D.L., Hackley, S., Lipp, O.V., Van Boxtel, A., 2005. Committee report: Guidelines for human startle eyeblink electromyographic studies. Psychophysiology 42, 1–15. https://doi.org/10.1111/j.1469-8986.2005.00271.x

Brainard, D.H., 1997. The Psychophysics Toolbox. Spat. Vis. 10, 433–436.

Brewin, C.R., 2001. A cognitive neuroscience account of posttraumatic stress disorder and its treatment. Behav. Res. Ther. 39, 373–393. https://doi.org/10.1016/S0005-7967(00)00087-5

Craske, M.G., 1999. Anxiety disorders: Psychological approaches to theory and treatment. Basic Books.

Craske, M.G., Mystkowski, J.L., 2006. Exposure Therapy and Extinction: Clinical Studies, in: Craske, M.G., Hermans, D., Vansteenwegen, D. (Eds.), Fear and Learning: From Basic Processes to Clinical Implications. American Psychological Association, Washington, DC, US, pp. 217–233. https://doi.org/10.1037/11474-011

Dunsmoor, J.E., Murphy, G.L., 2015. Categories, concepts, and conditioning: how humans generalize fear. Trends Cogn. Sci. 19, 73–77. https://doi.org/10.1016Zj.tics.2014.12.003

Ekman, P., 1976. Pictures of facial affect. Consult. Psychol. Press.

Fang, F., He, S., 2005. Cortical responses to invisible objects in the human dorsal and ventral pathways. Nat. Neurosci. 8, 1380–1385. https://doi.org/10.1038/nn1537

Golkar, A., Öhman, A., 2012. Fear extinction in humans: Effects of acquisition–extinction delay and masked stimulus presentations. Biol. Psychol. 91, 292–301. https://doi.org/10.1016/j.biopsycho.2012.07.007

Haerich, P., 1994. Startle Reflex Modification: Effects of Attention Vary With Emotional Valence. Psychol. Sci. 5, 407–410. https://doi.org/10.1111/j.1467-9280.1994.tb00294.x

Hamm, A.O., Vaitl, D., 1996. Affective learning: Awareness and aversion. Psychophysiology 33, 698–710. https://doi.org/10.1111/j.1469-8986.1996.tb02366.x

Hauner, K.K., Mineka, S., Voss, J.L., Paller, K.A., 2012. Exposure therapy triggers lasting reorganization of neural fear processing. Proc. Natl. Acad. Sci. 109, 9203–9208. https://doi.org/10.1073/pnas.1205242109

Hawk, L.W., Cook, E.W., 1997. Affective modulation of tactile startle. Psychophysiology 34, 23–31. https://doi.org/10.1111/j.1469-8986.1997.tb02412.x

J, W., Nadel, L., 1985. Stress-induced recovery of fears and phobias. Psychol. Rev. 92, 512–531. https://doi.org/10.1037/0033-295X.92A512

Kim, J.H., Richardson, R., 2010. New Findings on Extinction of Conditioned Fear Early in Development: Theoretical and Clinical Implications. Biol. Psychiatry, Posttraumatic Stress Disorder: Translational Neuroscience Perspectives on Gene-Environment Interactions 67, 297–303. https://doi.org/10.1016Zj.biopsych.2009.09.003

Kim, J.J., Baxter, M.G., 2001. Multiple brain-memory systems: the whole does not equal the sum of its parts. Trends Neurosci. 24, 324–330. https://doi.org/10.1016/S0166-2236(00)01818-X

Kindt, M., Soeter, M., 2013. Reconsolidation in a human fear conditioning study: A test of extinction as updating mechanism. Biol. Psychol., SI: Human Fear Conditioning 92, 43–50. https://doi.org/10.1016/j.biopsycho.2011.09.016

Koenigs, M., Huey, E.D., Raymont, V., Cheon, B., Solomon, J., Wassermann, E.M., Grafman, J., 2008. Focal brain damage protects against post-traumatic stress disorder in combat veterans. Nat. Neurosci. 11, 232–237. https://doi.org/10.1038/nn2032

Koizumi, A., Amano, K., Cortese, A., Shibata, K., Yoshida, W., Seymour, B., Kawato, M., Lau, H., 2016. Fear reduction without fear through reinforcement of neural activity that bypasses conscious exposure. Nat. Hum. Behav. 1, 0006. https://doi.org/10.1038/s41562-016-0006

Konarski, J.Z., Mcintyre, R.S., Soczynska, J.K., Kennedy, S.H., 2007. Neuroimaging Approaches in Mood Disorders: Technique and Clinical Implications. Ann. Clin. Psychiatry 19, 265–277. https://doi.org/10.3109/10401230701653435

Lang, P.J., Bradley, M.M., Cuthbert, B.N., 1990. Emotion, attention, and the startle reflex. Psychol. Rev. 97, 377–395.

Lang, P.J., Davis, M., Öhman, A., 2000. Fear and anxiety: animal models and human cognitive psychophysiology. J. Affect. Disord., Arousal in Anxiety 61, 137–159. https://doi.org/10.1016/S0165-0327(00)00343-8

Lapate, R.C., Rokers, B., Tromp, D.P.M., Orfali, N.S., Oler, J.A., Doran, S.T., Adluru, N., Alexander, A.L., Davidson, R.J., 2016. Awareness of Emotional Stimuli Determines the Behavioral Consequences of Amygdala Activation and Amygdala-Prefrontal Connectivity. Sci. Rep. 6. https://doi.org/10.1038/srep25826

LeDoux, J.E., 1993. Emotional memory systems in the brain. Behav. Brain Res. 58, 69–79. https://doi.org/10.1016/0166-4328(93)90091-4

LeDoux, J.E., 2015. Anxious: The Modern Mind in the Age of Anxiety by Joseph E LeDoux, book review. Oneworld Publications.

LeDoux, J.E., Brown, R., 2017. A higher-order theory of emotional consciousness. Proc. Natl. Acad. Sci. 114, E2016–E2025. https://doi.org/10.1073/pnas.1619316114

Lin, Z., He, S., 2009. Seeing the invisible: The scope and limits of unconscious processing in binocular rivalry. Prog. Neurobiol. 87, 195–211. https://doi.org/10.1016/j.pneurobio.2008.09.002

Lovibond, P.F., 2003. Causal beliefs and conditioned responses: Retrospective revaluation induced by experience and by instruction. J. Exp. Psychol. Learn. Mem. Cogn. 29, 97–106. https://doi.org/10.1037/0278-7393.29.1.97

McFadyen, J., Mermillod, M., Mattingley, J.B., Halász, V., Garrido, M.I., 2017. A Rapid Subcortical Amygdala Route for Faces Irrespective of Spatial Frequency and Emotion. J. Neurosci. 37, 3864–3874. https://doi.org/10.1523/JNEUROSCI.3525-16.2017

Méndez-Bértolo, C., Moratti, S., Toledano, R., Lopez-Sosa, F., Martínez-Alvarez, R., Mah, Y.H., Vuilleumier, P., Gil-Nagel, A., Strange, B.A., 2016. A fast pathway for fear in human amygdala. Nat. Neurosci. https://doi.org/10.1038/nn.4324

Milad, M.R., Quirk, G.J., 2012. Fear Extinction as a Model for Translational Neuroscience: Ten Years of Progress. Annu. Rev. Psychol. 63, 129–151. https://doi.org/10.1146/annurev.psych.121208.131631

Myers, K.M., Davis, M., 2002. Behavioral and Neural Analysis of Extinction. Neuron 36, 567–584. https://doi.org/10.1016/S0896-6273(02)01064-4

Oyarzún, J.P., Lopez-Barroso, D., Fuentemilla, L., Cucurell, D., Pedraza, C., Rodriguez-Fornells, A., de Diego-Balaguer R., 2012. Updating Fearful Memories with Extinction Training during Reconsolidation: A Human Study Using Auditory Aversive Stimuli. PLOS ONE 7, e38849. https://doi.org/10.1371/journal.pone.0038849

Pare, D., Duvarci, S., 2012. Amygdala microcircuits mediating fear expression and extinction. Curr. Opin. Neurobiol., Microcircuits 22, 717–723. https://doi.org/10.1016Zj.conb.2012.02.014

Pelli, D.G., 1997. The VideoToolbox software for visual psychophysics: transforming numbers into movies. Spat. Vis. 10, 437–442.

Phelps, E.A., Delgado, M.R., Nearing, K.I., LeDoux, J.E., 2004. Extinction Learning in Humans: Role of the Amygdala and vmPFC. Neuron 43, 897–905. https://doi.org/10.1016/j.neuron.2004.08.042

Porac, C., Coren, S., 1976. The dominant eye. Psychol. Bull. 83, 880.

Purkis, H.M., Lipp, O.V., 2001. Does Affective Learning Exist in the Absence of Contingency Awareness? Learn. Motiv. 32, 84–99. https://doi.org/10.1006/lmot.2000.1066

Raio, C.M., Carmel, D., Carrasco, M., Phelps, E.A., 2012. Nonconscious fear is quickly acquired but swiftly forgotten. Curr. Biol. CB 22, R477–479. https://doi.org/10.1016/j.cub.2012.04.023

Rescorla, R.A., 2004. Spontaneous Recovery. Learn. Mem. 11, 501–509. https://doi.org/10.1101/lm.77504

Rothbaum, B.O., Davis, M., 2003. Applying Learning Principles to the Treatment of Post-Trauma Reactions. Ann. N. Y. Acad. Sci. 1008, 112–121. https://doi.org/10.1196/annals.1301.012

Schiller, D., Kanen, J.W., LeDoux, J.E., Monfils, M.-H., Phelps, E.A., 2013. Extinction during reconsolidation of threat memory diminishes prefrontal cortex involvement. Proc. Natl. Acad. Sci. U. S. A. 110, 20040–20045. https://doi.org/10.1073/pnas.1320322110

Schiller, D., Monfils, M.-H., Raio, C.M., Johnson, D.C., LeDoux, J.E., Phelps, E.A., 2010. Preventing the return of fear in humans using reconsolidation update mechanisms. Nature 463, 49–53. https://doi.org/10.1038/nature08637

Sevenster, D., Beckers, T., Kindt, M., 2014. Fear conditioning of SCR but not the startle reflex requires conscious discrimination of threat and safety. Front. Behav. Neurosci. 8. https://doi.org/10.3389/fnbeh.2014.00032

Sevenster, D., Beckers, T., Kindt, M., 2012a. Retrieval per se is not sufficient to trigger reconsolidation of human fear memory. Neurobiol. Learn. Mem. 97, 338–345. https://doi.org/10.1016/j.nlm.2012.01.009

Sevenster, D., Beckers, T., Kindt, M., 2012b. Instructed extinction differentially affects the emotional and cognitive expression of associative fear memory. Psychophysiology 49, 1426–1435. https://doi.org/10.1111/j.1469-8986.2012.01450.x

Siegel, P., Warren, R., 2013a. The effect of very brief exposure on experienced fear after in vivo exposure. Cogn. Emot. 27, 1013–1022. https://doi.org/10.1080/02699931.2012.756803

Siegel, P., Warren, R., 2013b. Less is still more: maintenance of the very brief exposure effect 1 year later. Emot. Wash. DC 13, 338–344. https://doi.org/10.1037/a0030833

Siegel, P., Warren, R., Wang, Z., Yang, J., Cohen, D., Anderson, J.F., Murray, L., Peterson, B.S., 2017. Less is more: Neural activity during very brief and clearly visible exposure to phobic stimuli. Hum. Brain Mapp. 38, 2466–2481. https://doi.org/10.1002/hbm.23533

Siegel, P., Weinberger, J., 2009. Very brief exposure: The effects of unreportable stimuli on fearful behavior. Conscious. Cogn. 18, 939–951. https://doi.org/10.10167j.concog.2009.08.001

Soeter, M., Kindt, M., 2011. Disrupting reconsolidation: Pharmacological and behavioral manipulations. Learn. Mem. 18, 357–366. https://doi.org/10.1101/lm.2148511

Soeter, M., Kindt, M., 2010. Dissociating response systems: erasing fear from memory. Neurobiol. Learn. Mem. 94, 30–41. https://doi.org/10.1016Zj.nlm.2010.03.004

Sotres-Bayon, F., Cain, C.K., LeDoux, J.E., 2006. Brain Mechanisms of Fear Extinction: Historical Perspectives on the Contribution of Prefrontal Cortex. Biol. Psychiatry 60, 329–336. https://doi.org/10.1016/j.biopsych.2005.10.012

Spielberger, C.D., 1983. Manual for the State-Trait Anxiety Inventory STAI (Form Y) (“Self-Evaluation Questionnaire”).

Tsuchiya, N., Koch, C., 2005. Continuous flash suppression reduces negative afterimages. Nat. Neurosci. 8, 1096–1101. https://doi.org/10.1038/nn1500

Turk-Browne, N.B., Yi, D.-J., Chun, M.M., 2006. Linking Implicit and Explicit Memory: Common Encoding Factors and Shared Representations. Neuron 49, 917–927. https://doi.org/10.1016/j.neuron.2006.01.030

Wagnild, G.M., Young, H.M., 1993. Development and psychometric evaluation of the Resilience Scale. J. Nurs. Meas. 1, 165–178.

Warren, V.T., Anderson, K.M., Kwon, C., Bosshardt, L., Jovanovic, T., Bradley, B., Norrholm, S.D., 2014. Human fear extinction and return of fear using reconsolidation update mechanisms: The contribution of on-line expectancy ratings. Neurobiol. Learn. Mem., Extinction 113, 165–173. https://doi.org/10.1016/j.nlm.2013.10.014

Yang, E., Zald, D.H., Blake, R., 2007. Fearful expressions gain preferential access to awareness during continuous flash suppression. Emotion 7, 882–886. https://doi.org/10.1037/1528-3542.7.4.882

